# Development and validation of a real-time RT-PCR test for screening pepper and tomato seed lots for the presence of pospiviroids

**DOI:** 10.1101/2020.04.17.046508

**Authors:** Marleen Botermans, Johanna W. Roenhorst, Marinus Hooftman, Jacobus Th.J. Verhoeven, Eveline Metz, Esther J. van Veen, Bart P.J. Geraats, Mark Kemper, Debora C.M. Beugelsdijk, Harrie Koenraadt, Agata Jodlowska, Marcel Westenberg

## Abstract

Potato spindle tuber viroid and other pospiviroids can cause serious diseases in potato and tomato crops. Consequently, pospiviroids are regulated in several countries. Since seed transmission is considered as a pathway for the introduction and spread of pospiviroids, some countries demand for the testing of seed lots of solanaceous crops for the presence of pospiviroids. A real-time RT-PCR test, named PospiSense, was developed for testing pepper (*Capsicum annuum*) and tomato (*Solanum lycopersicum*) seeds for seven pospiviroid species known to occur naturally in these crops. The test consists of two multiplex reactions running in parallel, PospiSense 1 and PospiSense 2, that target Citrus exocortis viroid (CEVd), Columnea latent viroid (CLVd), pepper chat fruit viroid (PCFVd), potato spindle tuber viroid (PSTVd), tomato apical stunt viroid (TASVd), tomato chlorotic dwarf viroid (TCDVd) and tomato planta macho viroid (TPMVd, including the former Mexican papita viroid). Dahlia latent viroid (DLVd) is used as an internal isolation control. Validation of the test showed that for both pepper and tomato seeds the current requirements of a routine screening test are fulfilled, i.e. the ability to detect one infested seed in a sample of c.1000 seeds for each of these seven pospiviroids. Additionally, the Pospisense test performed well in an inter-laboratory comparison, which included two routine seed-testing laboratories, and as such provides a relatively easy alternative to the currently used tests.

## Introduction

Pospiviroids are single-stranded circular RNA molecules consisting of around 360 nucleotides. The genus *Pospiviroid* is in the family *Pospiviroidae*, with *Potato spindle tuber viroid* (PSTVd) being the type species. Most pospiviroids can infect a wide range of plant species, including many solanaceous ornamental and vegetable crops. Infected plants often remain symptomless, although PSTVd and some other pospiviroids may cause serious diseases in potato and tomato crops (1, 2). For this reason, many countries have implemented phytosanitary measures to prevent their introduction and spread.

Pospiviroids may spread by vegetative propagation, mechanical transmission, and to a lesser extent also by insects, pollen and seeds (3, 4).

The importance of seeds as a pathway for introduction and spread of pospiviroids in solanaceous fruit crops is still a matter of debate. This is due to the fact that both successful- and failed transmission from infested seeds to seedlings has been reported (5-8). Nevertheless, some countries require mandatory testing of pepper (*Capsicum annuum*) and tomato (*Solanum lycopersicum*) seed lots before import. Consequently, there is a need for reliable and cost-effective tests for screening pepper and tomato seed lots for PSTVd and other pospiviroids identified in these crops, i.e. Citrus exocortis viroid (CEVd), Columnea latent viroid (CLVd), pepper chat fruit viroid (PCFVd), tomato apical stunt viroid (TASVd), tomato chlorotic dwarf viroid (TCDVd) and tomato planta macho viroid (TPMVd, including the former Mexican papita viroid).

For detection of pospiviroids, several molecular tests are already available but they have their limitations regarding analytical specificity and sensitivity. The generic tests described by Botermans et al. (9), van Brunschot et al. (10), and Monger et al. (11) van were designed and validated for generic pospiviroid detection in leaf material, but are not sensitive enough for testing seed lots in which pospiviroid concentrations are generally lower. Other tests, such as the test described by Boonham et al. (12), are sensitive enough, but can only detect a limited number of species. Naktuinbouw (12-15) designed and validated a generic seed test, which is currently recommended by the International Seed Federation (16). This test consists of four parallel reactions that allow detection of one infested seed in a sample of c.1000 seeds for each of the seven pospiviroid species. A new test, therefore, should perform equally well and preferably reduces the number of reactions.

This paper describes the development and validation of a real-time RT-PCR test (PospiSense) for routine detection of the seven pospiviroid species in seeds of pepper and tomato. The test consists of two multiplex reactions running in parallel with a single internal isolation control, and provides an alternative to the currently used tests.

## Materials and methods

### Isolates used and confirmation of identity

Pospiviroid isolates and other pathogens used for test development and validation are presented in Table 1. The identity of the majority of pospiviroid species was confirmed by sequence analysis of the amplicons obtained by conventional RT-PCRs using different primer sets: Pospi1-FW/Pospi1-RE and VidRE/FW (17), Pospi2-FW/ Pospi2-RE (18), the primers described by Shamloul et al. (19) and AP-FW1/RE2 (20). Because of the lower analytical sensitivity of primers VidRE/FW (17) and the primers described by Spieker (21) no amplicon or sequence data were obtained for CLVd isolates from samples 6184939, PPS013 and PPS055. Therefore its presence and identity was verified by a CLVd-specific real-time RT-PCR published by Monger et al. (11). Amplicons were bi-directionally sequenced as described by Van de Vossenberg and Van der Straten (22). The identity of the virus isolates was confirmed by ELISA or sequencing. The identity of *Clavibacter michiganensis* subsp. *michiganensis* was confirmed by real-time PCR and a pathogenicity test.

**Table 1.**
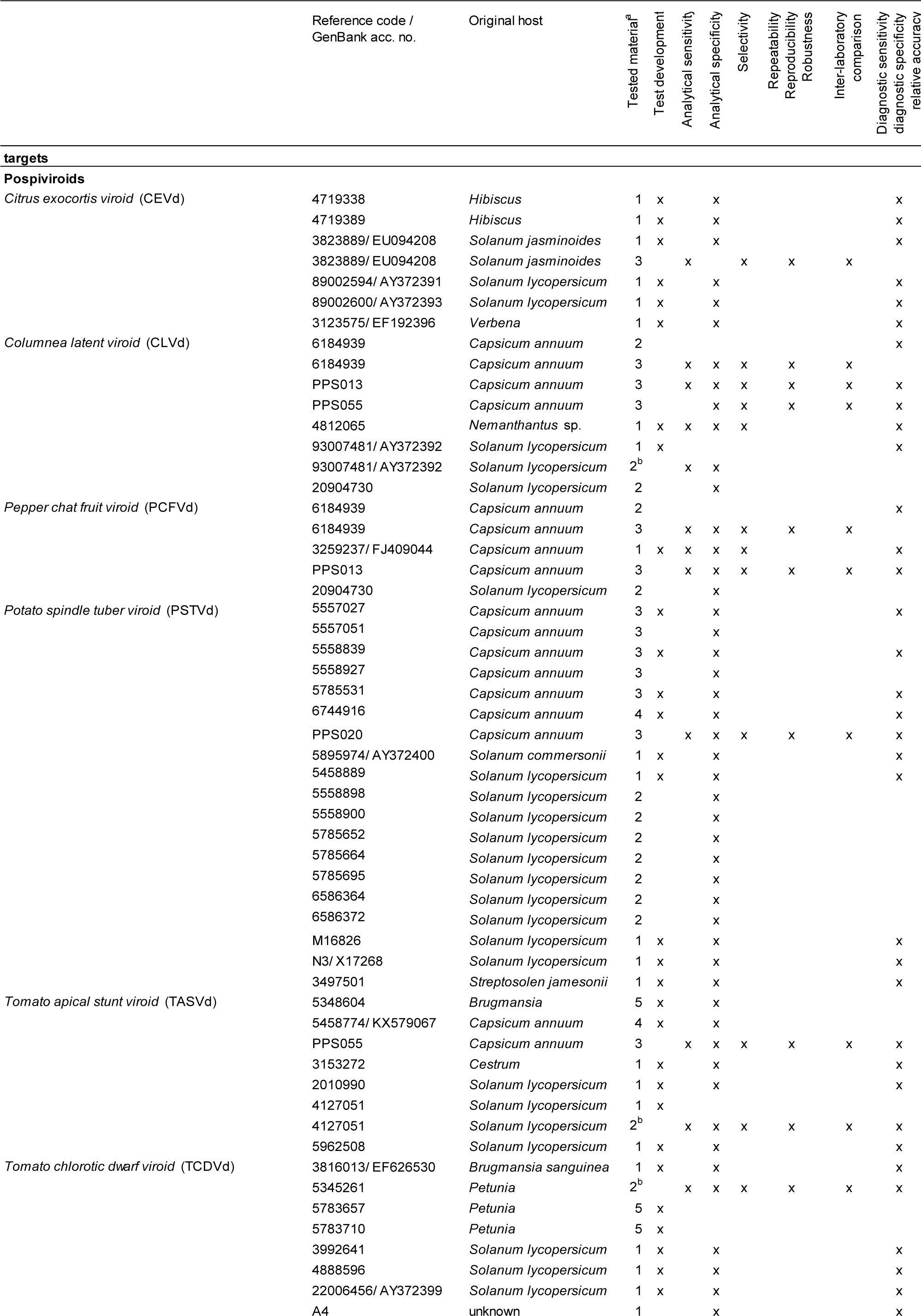
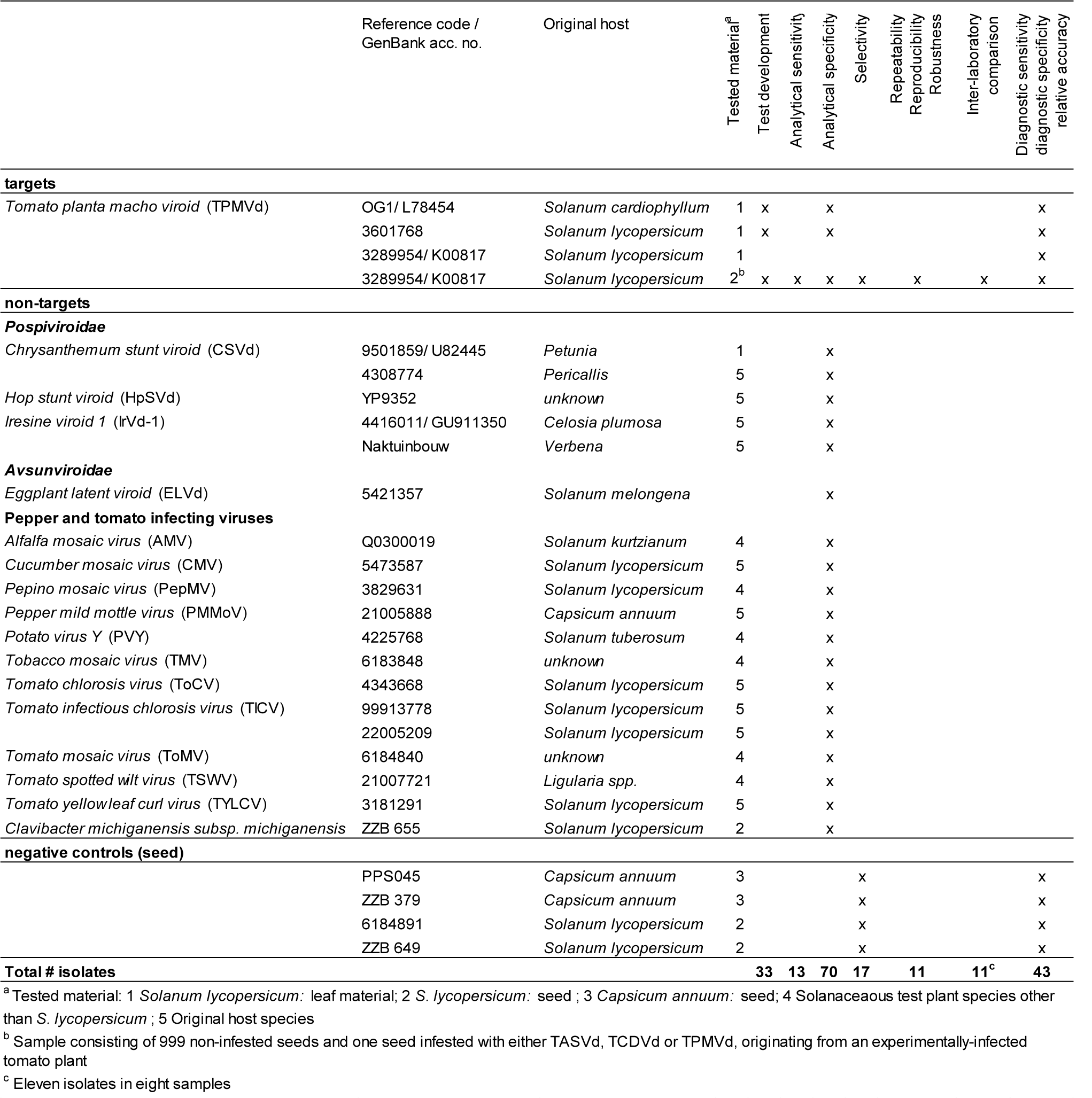
Overview of isolates (targets and non-targets) and control material used in this study.

### Test development

Complete genome sequences of the target species (CEVd, CLVd, PCFVd, PSTVd, TASVd, TCDVd and TPMVd) were retrieved from NCBI GenBank and the sequence database of the National Plant Protection Organization of the Netherlands. For all seven pospiviroids sequences of over 130 isolates covering the intra-species variation were selected. Sequences were aligned with the MAFFT alignment tool (23) in Geneious R8 (Biomatters) and manually adjusted. To minimise the number of reactions, primer and probe design focused on the conserved regions shared by different combinations of the seven species. Potentially suitable sites for primers and probes were visually identified and the oligonucleotide design further optimised using PrimerExpress 3 (Thermo Fischer Scientific). Primers and probes were tested in different combinations together with published primers and probes for CLVd (11), resulting in the selection of primers and probes listed in Table 2. Since the selected primers and probes could not be combined in one single reaction without losing (analytical) sensitivity, the final design of the PospiSense test consisted of two reactions run in parallel, named PospiSense 1 and PospiSense 2.

**Table 2.**
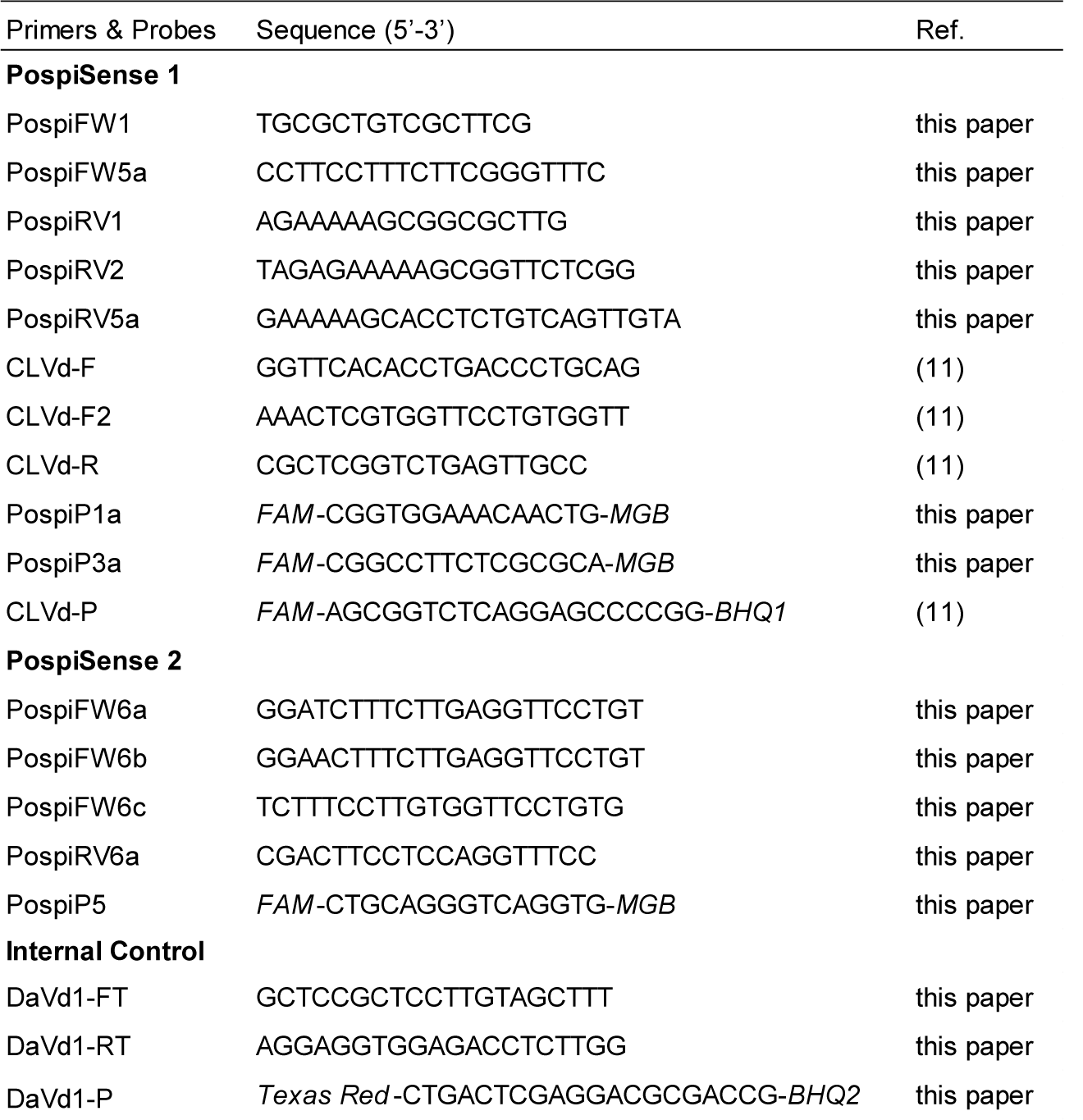
Primers and probes sequences of PospiSense test.

In both reactions, dahlia latent viroid (DLVd; genus *Hostuviroid*) was included as an (exogenous) internal isolation control, which was detected by using the primers and probe (Table 2).

### Sample preparation and RNA extraction

#### Seeds

Samples of tomato seeds consisted of c. 3000 seeds, which were divided in three subsamples of c.1000 seeds for testing, according to standard procedures used by seed testing laboratories in Europe (16).

Seeds were processed by either using a Geno/Grinder (SPex SamplePrep P) or a BagMixer 100 (Interscience), depending on the laboratory’s preference. When using the Geno/Grinder (dry processing) 3x 1000 tomato seeds were transferred to a 50 ml tube containing a 14 mm steel ball. Tubes were put upside down and seeds ground at 1500 rpm for 4 min (at least 95% seeds crushed). After grinding, 20 ml GH+ extraction buffer ((9); modified by (24)) which included the DLVd spike, was added to each tube. The DLVd-spike stock of 1 g DLVd-infected leaf material homogenised in 10 ml GH+ buffer was used at a dilution of approximately 10^−4^ to achieve a Cq value of about 28. To obtain homogenous solutions the tubes were shaken manually. When using the BagMixer (wet processing) subsamples of 1000 tomato seeds were transferred to a grinding bag (BagPage 100ml (Interscience) and soaked in 20 ml GH+ buffer spiked with DLVd at room temperature for 30-60 min and subsequently blended for 1.5 min at position 4.

For pepper seeds the same procedure was followed except that the subsamples of c.1000 seeds were subdivided into 2×500 seeds before grinding with a Geno/Grinder for 7 min. After grinding 10 ml DLVd-spiked GH+ extraction buffer was added to each of the six tubes, followed by combining and mixing of the contents of the two tubes of a subsample before further processing.

For RNA extraction, 1 ml of the seed homogenate was transferred into a 1.5 ml tube and 30 μl of 5M Dithiothreitol was added, followed by incubation in a thermoshaker at 850 rpm, 65°C for 15 min. The tubes were centrifuged at 16,000 g for 10 min. For manual RNA extraction using the RNeasy plant mini kit (Qiagen), 750 μl of the supernatant was used following the manufacturer’s instructions. For large-scale RNA extraction on a Kingfisher KF96 system (Thermo Fisher Scientific), using the Sbeadex Maxi Plant kit (LGC), 250 μl of supernatant was transferred to a binding plate containing 600 μl of binding buffer and 50 μl Sbeadex particle suspension, following the manufacturer’s instructions.

#### Leaves

Sample preparation and RNA extraction from leaf material (Table 1) was performed according to Botermans et al. (9).

### PospiSense real-time RT-PCR

Table 2 lists the primer and probe sequences used for the PospiSense test. The PospiSense 1 reaction contained: 1x UltraPlex 1-Step ToughMix (Quanta Biosciences), 0.3 µM of each PospiSense 1 and internal control primer, 0.1 µM of TaqMan probe PospiP1a, PospiP3a, CLVd-P, 0.2 µM of internal control TaqMan probe DaVd1-P, 2 µl RNA template and molecular grade water to a final volume of 20 µl. The PospiSense 2 reaction included: 1x UltraPlex 1-Step ToughMix (Quanta Biosciences), 0.3 µM of each PospiSense 2 and internal control primer, 0.1 µM of TaqMan probe PospiP5, 0.2 µM of internal control TaqMan probe DaVd1-P, 2 µl RNA template and molecular grade water to a final volume of 20 µl. Both reactions used real-time RT-PCR: 10 min 50°C, 3 min 95°C, followed by 40 cycles 10 s 95°C and 1 min 60°C. Real-time RT-PCRs were carried out in 96-well plates on a Bio-Rad CFX96™ Real-Time PCR system (Bio-Rad Laboratories,) or a QuantStudio™ 6 Flex Real-Time PCR System (Thermo Fisher Scientific). After verification of controls, a test result was considered positive if an exponential amplification curve was produced for either the PospiSense 1 and/or the PospiSense 2 reaction.

## Results

### Test development and validation

Table 2 shows the primers and probes that were selected for further validation, based on the results of the initial tests. To determine whether the PospiSense test is suitable for routine testing of seed lots, the following performance characteristics were determined: analytical sensitivity, analytical specificity, selectivity, repeatability and reproducibility, according to the EPPO standard PM7/98 version 4 (25). In addition, the PospiSense test was compared with the currently most-commonly used test (14, 15) for diagnostic sensitivity, diagnostic specificity and relative accuracy (25). Table 1 indicates the isolates used to determine each of the performance characteristics.

#### Analytical sensitivity

To determine the analytical sensitivity, RNA-extracts of pepper or tomato seeds naturally infested by CLVd, PSTVd or TASVd (one isolate each) were diluted in duplicate in RNA-extracts of non-infested seeds. Testing of RNA extracts of decimal dilutions revealed that these three pospiviroids showed 100% detection up to 1000, 10.000 and 100 times dilution respectively. Furthermore, testing samples consisting of one tomato seed infested by either CLVd, TASVd, TCDVd or TPMVd, and 999 non-infested seeds, produced consistent positive results. In comparison to the Naktuinbouw test, the PospiSense test appeared less sensitive for detection of CEVd and TASVd, (difference for CEVd ΔCq= 6.1 SD=3.2 n=6 and TASVd ΔCq= 4.0 SD=1.4 n=5, based on average values of both leaf and seed samples in a range of concentrations). Nevertheless, the PospiSense test meets the requirements of detecting one infested seed in a sample of c.1000 seeds.

#### Analytical specificity

The analytical specificity was determined by testing infected leaf and infested seed samples by target and non-target species (see Table 1). The PospiSense test gave positive results for all 51 tested isolates of the seven target pospiviroids, i.e. CEVd (6), CLVd (6), PCFVd (4), PSTVd (19), TASVd (7), TCDVd (6) and TPMVd (3), thus showing coverage of the intra-species variability (inclusivity). For 12 non-targets (exclusivity), no cross-reactions were observed, i.e. for hop stunt viroid (hostuviroid), and most common pepper- and tomato-infecting viruses, i.e. alfalfa mosaic virus, cucumber mosaic virus, pepper mild mottle virus, pepino mosaic virus, potato virus Y, tobacco mosaic virus, tomato chlorosis virus, tomato mosaic virus, tomato spotted wilt virus and tomato yellow leaf curl virus. In addition, no cross-reactions were found for the bacterium *Clavibacter michiganensis* subsp. *michiganensis*. For four non-targets, i.e. Chrysanthemum stunt viroid (CSVd), eggplant latent viroid (genus *Elaviroid*) and Iresine viroid 1 (IrVd-1), cross-reactions (Cq= 27-37) were observed when present in high concentrations. Of these viroid species, however, no natural infections in pepper and tomato have been reported. In addition, one out of two isolates of tomato infectious chlorosis virus showed some cross-reactivity when present in high concentration, which is not likely for seeds. Moreover during confirmatory testing, false positives will be revealed and abolished. Therefore, the observed cross-reactions will not hamper the application of the PospiSense test for screening seed lots.

#### Selectivity

To determine the effect of the matrix, RNA extracts of seeds containing RNA of each of the target species were diluted in either RNA-extracts of non-infested seeds or water (Table 1). Test results of serial dilutions of the RNA extracts were compared. For both pepper and tomato seeds only minor differences were observed, i.e. average Δ Cq water: pepper seed= 0.4 and average Δ Cq water: tomato seed= 0.6. Regarding selectivity, therefore, it was concluded that no apparent matrix effects occurred in both pepper and tomato seeds.

#### Repeatability and Reproducibility

Repeatability and reproducibility were determined by analysing sub-samples of pepper and tomato seeds under the same experimental conditions (technical replicates) and under different experimental conditions (date, operator, apparatus, *etc*.), including an inter-laboratory comparison. Eight samples of pepper and tomato seeds infested by the seven relevant pospiviroids (11 isolates) were selected. The RNA extracts of these samples with medium to low relative infestation rates (Cq values of targets varying between 20 and 32) were sub-sampled and tested by the three participating laboratories, including two routine seed-testing laboratories. Qualitative interpretation of the resulting data showed concordance for all (sub-) samples, both within and between laboratories, and irrespective of variation in experimental conditions (Table 3a,b). Repeatability and reproducibility were 100%, further demonstrating the robustness of the PospiSense test.

**Table 3a.** Results (Cq values) of the repeatability and reproducibility experiments in intra- and inter-laboratory setting (PospiSense 1)

| Isolate | Matrix (seed) | PospiSense 1, Test Moment 1-6<sup>a</sup> |  |  |  |  |  |
| --- | --- | --- | --- | --- | --- | --- | --- |
|  |  | NPPO Lab |  |  |  | Lab 1 | Lab 2 |
|  |  | 1 | 2 | 3 | 4 | 5 | 6 |
| PSTVd PPS020 | Pepper | 22/22 | 22 |  |  | 23 |  |
| TASVd + CLVd PPS055 | Pepper | 25/25 | 25 |  |  | 26 |  |
| PCFVd + CLVd PPS013 | Pepper | 24 | 25/24 |  |  | 28(VIC)<sup>d</sup> |  |
| TCDVd 5345261<sup>b</sup> | Tomato | 24 | 24/24 |  |  |  | 23 |
| TPMVd 3289954<sup>b</sup> | Tomato |  |  | 29/30 | 30 |  | 28 |
| TASVd 4127051<sup>b</sup> | Tomato |  |  | ND/ND | ND |  | ND |
| PCFVd + CLVd 6184939 | Pepper |  |  | 23 | 22/22 |  | 22 |
| CEVd 3823889<sup>c</sup> | Pepper |  |  | ND | ND/ND |  | ND |

**Table 3b.** Results (Cq values) of the repeatability and reproducibility experiments in intra- and inter-laboratory setting (PospiSense 2)

| Isolate | Matrix (seed) | PospiSense 2, Test Moment 1-6<sup>a</sup> |  |  |  |  |  |
| --- | --- | --- | --- | --- | --- | --- | --- |
|  |  | NPPO Lab |  |  |  | Lab 1 | Lab 2 |
|  |  | 1 | 2 | 3 | 4 | 5 | 6 |
| PSTVd PPS020 | Pepper | ND/ND | ND |  |  | ND |  |
| TASVd + CLVd PPS055 | Pepper | 25/25 | 25 |  |  | 25 |  |
| PCFVd + CLVd PPS013 | Pepper | 37 | 37/37 |  |  | 38 |  |
| TCDVd 5345261<sup>b</sup> | Tomato | ND | ND/ND |  |  |  | ND |
| TPMVd 3289954<sup>b</sup> | Tomato |  |  | ND/ND | ND |  | ND |
| TASVd 4127051<sup>b</sup> | Tomato |  |  | 32/32 | 32 |  | 31 |
| PCFVd + CLVd 6184939 | Pepper |  |  | 37 | 37/37 |  | 37 |
| CEVd 3823889<sup>c</sup> | Pepper |  |  | 20 | 20/20 |  | 19 |

<sup>a</sup> 1-6 = Moment at which test is performed by a different operator
<sup>b</sup> RNA extract from sample consisting of 999 non-infested seeds and one seed infested with TASVd, TCDVd or TPMVd, originating from an artificially infected tomato plant.
<sup>c</sup> RNA extract from seed spiked with CEVd
<sup>d</sup> PCFVd specific probe was accidentally labeled with VIC instead of FAM fluorophore, explaining slightly different cq values
ND = not detected (negative test result)
Test results of internal control were positive (data not shown)

#### Diagnostic sensitivity, diagnostic specificity and relative accuracy

To determine the relative accuracy of the PospiSense test, test results were compared with the results obtained with the with the most-commonly used pospiviroid seed test of Naktuinbouw (2017a,b,c). In total, 43 samples including both infested and non-infested seed samples were tested with both tests. Positive and negative results were compared qualitatively. The PospiSense and the Naktuinbouw test both diagnosed the same number of positive (n=39; Naktuinbouw test Cq 12-31, PospiSense test Cq 10-34) and negative (n=4) results. Consequently, diagnostic sensitivity, diagnostic specificity and relative accuracy were all 100% in comparison with the Naktuinbouw test.

## Discussion

The newly developed PospiSense test has been shown to fulfil the requirements for routine testing of pepper and tomato seed samples for the seven pospiviroid species known to occur naturally in these crops. The validation data showed that for CLVd, TASVd, TCDVd and TPMVd the test allows detection of at least one infested seed in a sample of 1000 seeds. For PSTVd similar results were obtained for single seeds from a naturally infested seed lot. For CEVd no infested seed samples were available, and for PCFVd only seed samples co-infested with CLVd. Therefore, the analytical sensitivity could not be experimentally determined. However, the results are expected to be similar, because these pospiviroids are likely to share both physical and biological characteristics with other members in the genus. Moreover, regarding test performance, the analytical sensitivity for CEVd and PCFVd for leaf material is within the same range as the other pospiviroids. In addition, the results of the wide range of targets and non-targets tested, as well as the absence of matrix effects, showed its suitability for screening both pepper and tomato seeds. A 100% repeatability and reproducibility were obtained during validation and inter-laboratory comparison, both further demonstrating the robustness of the PospiSense test.

For routine screening of seed lots, the PospiSense test offers some improvements in comparison with the currently used real-time RT-PCR pospiviroid tests. Firstly, the test is more sensitive than the other (semi-) generic pospiviroid tests as described by Monger et al. (11) and Botermans et al. (9), which both lack the sensitivity needed for reliable seed testing. Secondly, the PospiSense test is less complex than the pospiviroid seed test of Naktuinbouw (14, 15) and its performance characteristics generally comparable, although the analytical sensitivity of the PospiSense is slightly lower for CEVd and TASVd. In addition, the comparison of both tests showed a 100% agreement. However, in comparison to the Naktuinbouw test, the PospiSense test consists of two instead of four parallel reactions and uses only one internal control (DLVd) and one fluorophore. In both reactions, DLVd is spiked as internal isolation control. This control appeared a more consistent control for seed testing than the host-derived nad5, which often produces variable Cq values due to differences in cell physiology. The characteristics of the DLVd control are similar to the targets and its secondary structure is likely to prevent it from degradation by RNases. Another factor contributing to the lesser complexity of the PospiSense test is the choice of using the same fluorophore for all target species, as it makes the interpretation of test results easier. There is little chance of confusing results caused by cross-reactions between different primers and probes and/or the presence of more than one pospiviroid species in a sample. Nevertheless, it is possible to include additional fluorophores if discrimination among species at the screening stage is desirable.

The PospiSense test has been developed for efficient testing of seeds by combining the detection of seven pospiviroid species. This implies that in the case of a positive result, at least one pospiviroid species could be present and additional tests are needed for the identification of the species. Specific real-time RT-PCR tests have been developed to detect CEVd, CLVd, TASVd (26), and PCFVd (14, 15). For the closely related species PSTVd, TCDVd and TPMVd, the real-time RT-PCR test described by Boonham et al. (2004) can be used for confirmation, but by detecting all these three pospiviroids (except for one TPMVd isolate), the test is not able to distinguish between these species. Consequently, these three species can only be distinguished and identified by sequencing the amplicons obtained by conventional RT-PCR. Furthermore, it should be noted that for confirmation, a different test, preferably targeting a different region of the genome, should be used. However, the identification of pospiviroids in seed lots is not always easy, since viroid concentrations are generally low. Identification has even become more challenging because of the increased sensitivity of the recently developed real-time RT-PCR tests, including the PospiSense test described in this paper. According to the International Committee on Taxonomy of Viruses (27), the identification of viroids should be based on the analysis of their complete genome. Complete sequences, however, are still difficult or impossible to obtain from seed samples with low viroid levels, because conventional RT-PCR tests lack the required sensitivity to produce full-length amplicons. In addition, in comparison to real-time RT-PCR tests, conventional RT-PCRs are generally more prone to inhibition by matrix components. This means that seed treatments might have more impact on the analytical sensitivity of the conventional RT-PCR tests. For the identification of pospiviroid species in seed samples, the primer set Pospi1-FW/Pospi1-RE described by Verhoeven et al. (17) appeared most suitable due to its relatively high analytical sensitivity (28) When combining this test with the primer set Pospi2-FW/ Pospi2-RE (18), complete genome sequences of all known pospiviroids (except for CLVd) can be obtained, since these two primer pairs anneal at the same loci but in opposite polarity. However, often tailor-made solutions are needed for the confirmation and identification of pospiviroids in seed samples, e.g. concentration methods (29) nested-RT-PCR, or pooling of PCR-products for further testing. In conclusion, the performance of the PospiSense test, combined with the need of only two parallel reactions and a limited number of probes, shows its perspectives as an alternative test for screening seed lots of solanaceous species.

## Acknowledgements

We would like to thank Adrian Fox and colleagues from Fera Science Ltd (York, UK) and Ricardo Flores from Instituto de Biología Molecular y Celular de Plantas (UPV-CSIC) (Valencia, Spain) for providing isolates, and Joris Voogd, Tim Warbroek and Maureen Bruil for technical assistance.

